# Assessing the impact of binary land cover variables on species distribution models: A North American study on water birds

**DOI:** 10.1101/2023.07.31.551237

**Authors:** Lukas Gabor, Jeremy Cohen, Walter Jetz

## Abstract

**Aim:** Species distribution models (SDMs) are an important tool for predicting species occurrences in geographic space and for understanding the drivers of these occurrences. An effect of environmental variable selection on SDM outcomes has been noted, but how the treatment of variables influences models, including model performance and predicted range area, remains largely unclear. For example, although landcover variables included in SDMs in the form of proportions, or relative cover, recent findings suggest that for species associated with uncommon habitats the simple presence or absence of a landcover feature is most informative. Here we investigate the generality of this hypothesis and determine which representation of environmental features produces the best-performing models and how this affects range area estimates. Finally, we document how outcomes are modulated by spatial grain size, which is known to influence model performance and estimated range area.

**Location:** North America

**Methods:** We fit species distribution models (via Random Forest) for 57 water bird species using proportional and binary estimates of water cover in a grid cell using occurrence data from the eBird citizen science initiative. We evaluated four different thresholds of feature prevalence (land cover representations) within the cell (1%, 10%, 20% or 50%) and fit models across both breeding and non-breeding seasons and multiple grain sizes (1, 5, 10, and 50 km cell lengths).

**Results:** Model performance was not significantly affected by the type of land cover representation. However, when the models were fitted using binary variables, the model-assessed importance of water bodies significantly decreased, especially at coarse grain sizes. In this binary variable-case, models relied more on other land cover variables, and over-or under-predicted the species range by 5-30%. In some cases, differences up to 70% in predicted species ranges were observed.

**Main conclusions:** Methods for summarizing landcover features are often an afterthought in species distribution modelling. Inaccurate range areas resulting from treatment of landcover features as binary or proportional could lead to the prioritization of conservation efforts in areas where the species do not occur or cause the importance of crucial habitats to be missed. Importantly, our results suggest that at finer grain sizes, binary variables might be more useful for accurately measuring species distributions. For studies using relatively coarse grain sizes, we recommend fitting models with proportional land cover variables.

## Introduction

The relationship between species and their environment is fundamental to ecology (Humbold von and Bonpland 1807, Hairston 1959, Neyman and Scott 1959, MacArthur 1960) and gaining new importance in the face of ongoing global change (Butchart et al. 2010, Barnosky et al. 2011, Carlson et al. 2022). Species distribution models (SDMs) are a widely used tool in ecology that enable the description of species-environment relationships and, consequently, the prediction of species-habitat relationships in environmental space (see, for example, Václavík et al. 2012, Cord et al. 2013, Cohen et al. 2016, Marzialetti et al. 2019, Ellis-Soto et al. 2021, Lindegren et al. 2022, Cogliati et al. 2023). Even though species distribution models are now commonly adopted, scientists still face challenges related to the quality and type of the input data (both species and environmental), which can significantly impact the fitted models (Araújo et al. 2019, Moudrý et al. 2019, Gábor et al. 2020, Smith and Santos 2020, Bazzichetto et al. 2022, Zarzo-Arias et al. 2022, Moudrý et al. 2023, Smith et al. 2023). While numerous studies have shown species distribution models to be greatly affected by the environmental features that are included (e.g., Cord et al. 2014, McCluskey et al. 2018, Howard et al. 2020), recent work suggests that the summarization of these features as either proportional or binary also has potential to affect the model quality, the predicted area of occurrence, and the variables that the model associates with species occurrence (Gábor et al. 2022a).

Land cover features, which typically describe the available habitat within a spatial unit, are commonly included as environmental variables in SDMs. These variables are often represented by the area or proportion of a specific land cover type within the study area (e.g., Lecours et al. 2020, Rose et al. 2020, Coppée et al. 2022, Koma et al. 2022, Peng et al. 2022). In a recent study, Gábor et al. (2022a) demonstrated that for species specializing in locally rare habitats, such as water bodies in central Europe, the amount of habitat within a spatial unit is sometimes less important than the presence of the habitat itself. As a result, they recommended using binary land-cover variables to develop SDMs in regions with such habitats. They further suggested extending this approach to species that specialize in prevalent habitats, such as forests in central Europe, based on the concept of critical habitat area (Andrén 1994, Fahrig 2001, Melo et al. 2018). This concept hypothesized that there is a threshold in habitat amount below which a species cannot survive – e.g., loons (order Gaviiformes) are physiologically constrained from foraging on land and require water habitat for survival (see Figure 1). Thus, the authors proposed to find and set a threshold above which the species is expected to persist (e.g., 20%) and to use this threshold to derive binary land-cover variables. However, the authors highlighted that further research is needed to confirm this hypothesis, especially using various spatial scales or grain sizes (i.e., resolution of environmental variables).

**Figure 1.**
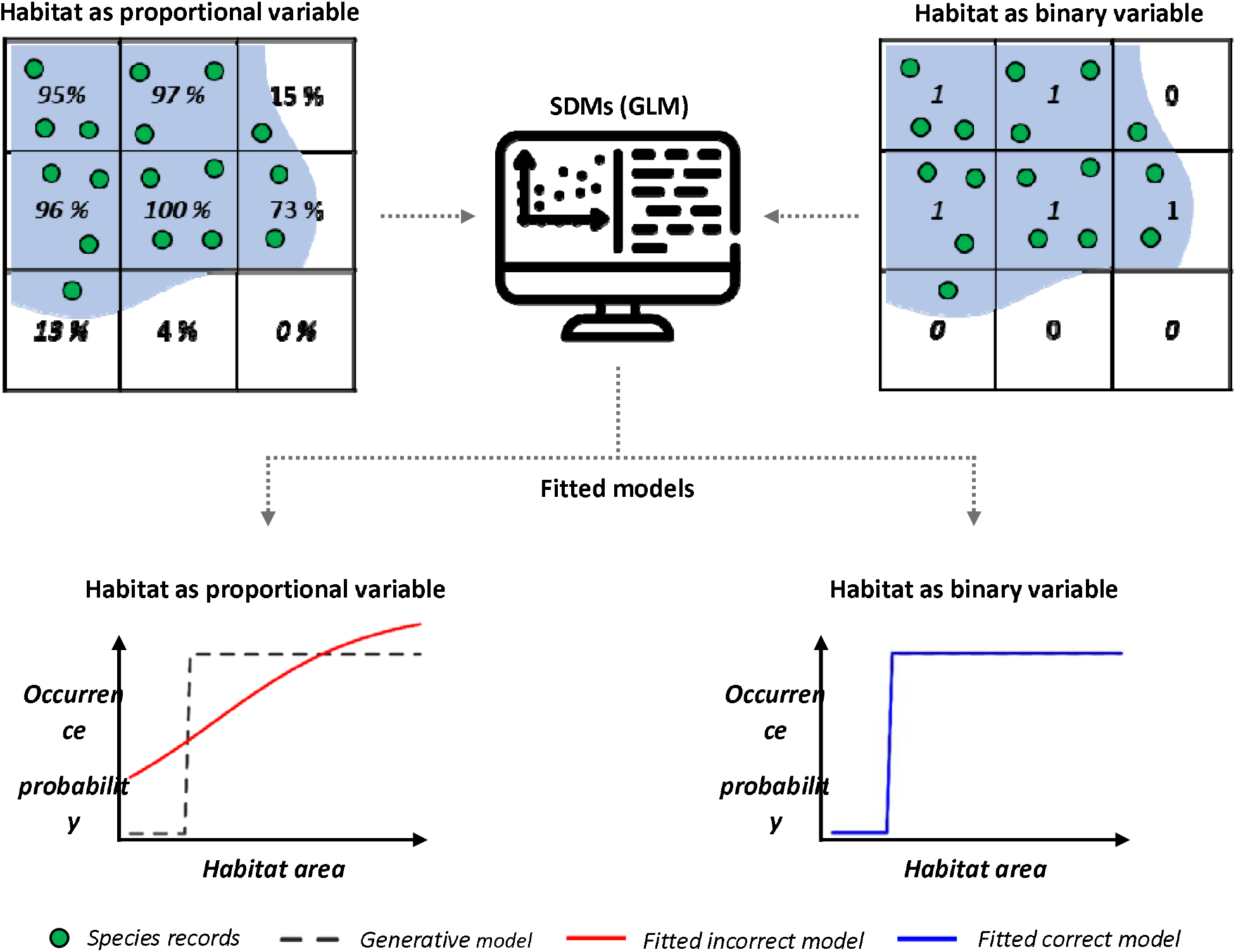
Graphical rationale for the binary variables’ hypothesis. We modelled species occurrence probability as a step function (generative model, black dashed line) to generate 100 presences/absences of a species that needs at least 20% of landcover within a spatial unit, drawn from a Bernoulli distribution with species occurrence probability as a parameter. Then, we fitted species distribution models (GLM) with either proportional (incorrect model, red line) and binary coverage (correct model, blue line), as variable.

The scale dependence of species-environment relationships is increasingly noted (Mertes and Jetz 2018, Luebert et al. 2022, Lu & Jetz 2023). Nowadays, species distribution models are developed at a wide range of spatial scales and grain sizes (i.e., the resolution of environmental variables), from a few meters (e.g., Bazzichetto et al. 2018, Lecours et al. 2020, Casanelles-Abella et al. 2022, Stark and Fridley 2022) to many kilometres (e.g., Kleisner et al. 2017, Norberg et al. 2019, Zarzo-Arias et al. 2022). The issue of grain size can also be critical for determining the applicability of land-cover binary variables for prevalent habitats. Fine resolution data offers a wider range of threshold options to derive binary land cover variables. Conversely, larger grain size requires a lower threshold. For instance, a threshold of up to 50% might be meaningful when using a 1 x 1 km grain size to derive a binary variable for water bodies. However, this threshold would not be appropriate when using a grain size of 20 x 20 km, as it would indicate only a small number of water bodies. Consequently, fitted models for water bird species could inappropriately indicate that the most important variable is the land cover surrounding water bodies instead of the water bodies themselves. Although such models can still achieve very good performance, there may be unintended consequences for their ecological interpretation, predicted species distribution, and conservation actions aimed to mitigate the decline in biodiversity due to climate change across local to global scales.

This study examines how the representation of a land cover variable affects model performance, range area estimation, and the importance of other habitat characteristics to the model. To accomplish this, we individually fit species distribution models for 57 water bird species in North America for both breeding and non-breeding seasons using eBird citizen science data. We fit models across both seasons because many of these species occupy highly distinct ranges across each season and some are more dependent on water during the breeding season, when offspring cannot yet fly (Zuckerberg et al. 2016). The eBird dataset consists of extensive and often spatially detailed occurrence data, making it ideal to model species distributions down to fine spatial scales across large spatial extents and across multiple times of year. For each species, we fit models with 1) water cover summarized proportionally (i.e., amount within the spatial unit) or at binary thresholds of 1%, 10%, 20% and 50%, and 2) environmental covariates summarized at a grain size of 1 x 1 km, 5 x 5 km, 10 x 10 km and 50 x 50 km.

Specifically, we address the following questions: (a) Do models built with binary landcover variables perform better than those built with proportional variables, and if so, what is the role of chosen threshold to derive binary variables? (b) How does the choice of land cover summary (i.e., binary versus proportional) affect the choice of landcover types that the model associates with the species? (c) How will this choice affect the prediction of suitable land area? (d) What role does the grain size of land cover variables play in this process? (e) Is the inclusion of proportional or binary landcover type most important during the breeding season, when species are more closely tied to specific habitat types?

## Methods

### Modeling region and species selection

We modelled species distributions across the North American continent, a box with dimensions (179.99°W, 42.68°W, 10.53°S, and 87.11°N) that included Canada, the United States, and the majority of Central America. Although we only modelled species native to the United States and Canada and only present biodiversity estimates for this region, modeling each species’ entire continental range was required to ensure accurate predictions.

Based on American Birding Association birding codes (2008) updated to the Clements bird taxonomy as of 2021, we compiled a list of 197 water-associated native bird species that breed or overwinter in the United States and Canada each year (Clements, 2007). These codes are widely used to distinguish regularly occurring species (code 1 or 2 species, which we use) from vagrants occurring irregularly (code 3+). We excluded species with almost entirely marine ranges or with insufficient data points and those that have records only for one season (i.e., breeding, non-breeding). We also avoided modeling Hawaiian endemics because these species were restricted to islands that were smaller than some of our spatial grain sizes.

### Data acquisition

We gathered data from eBird, a global citizen science initiative in which users submit checklists containing bird observations that has become widely used for understanding species distributions at high resolution (Sullivan et al., 2014). Users can indicate whether or not all observed species were recorded on the checklist (“complete” checklists), allowing for absence inference and presence-absence modeling. Users also indicate the level of effort involved in each observation by providing the distance travelled, time spent birding, and number of observers (hereafter, effort indicators).

R 4.1.0 was used to complete all data compilation, analyses, and visualizations (R Core Team, 2021). We compiled all eBird data for all species separately during the breeding season (June-August; June-July for shorebirds or Charadriiformes, which migrate early) and non-breeding season (December-February). We removed subspecies information from all checklists and summarized all data at the species level. Following established eBird data modeling protocols, we initially applied several filters to the data to reduce bias and improve data quality. (Johnston et al., 2019; Kelling et al., 2018). First, we eliminated checklists with extremely long durations (> 3 hours), large numbers of observers (>5), or protocols other than “stationary” or “traveling,” as these are incomparable with the majority of eBird’s data. Second, to reduce observational positional error, we eliminated checklists covering more than 1 km because they are likely to result in greater spatial uncertainty. Third, data prior to 2004 were removed because there is insufficient data from earlier years to adequately control for long-term temporal trends. Finally, data from users with fewer than five contributions were removed to reduce bias (e.g., false absence) caused by inexperienced birders.

Before modeling, the data was further filtered at the species level. Checklists were limited to those falling within a 200 km buffer of the species’ seasonal expert range boundary to limit overprediction outside of the species’ range extent (via Cornell’s spatial boundaries - https://ebird.org/science/status-and-trends/download-data, accessed July 2021). When modeling shorebirds (order Charadriiformes), we excluded August data because many species are already migrating long distances by this time. We limited checklists to one per 5 km grid cell per week to reduce site selection and temporal bias in data collection. To reduce the imbalance between presence and absence points, we repeated this filtering for checklists where the species occurs and does not occur. Coordinates, polygons and grids used in the study operated under a conical equal area projection. For spatial geoprocessing, the raster (Hijmans et al., 2015), rgdal (Bivand et al., 2015), and sf (Pebesma, 2018) packages were used. We ended up modeling 57 species (see Table A1).

### Landcover data

For the following categories, our landcover suite of covariates included percent landcover within a buffer (with size corresponding to modeling grain size) corresponding to the year the checklist was recorded: mixed forest, mosaic, shrubland, grassland, lichens/mosses, sparse, flooded/freshwater, flooded/saltwater, flooded/shrub, urban, barren, and ice (from European Space Agency – Climate Change Initiative; “ESA. Land Cover CCI Product User Guide Version 2. Technical Report.,” 2017).

To examine how water cover influences model performance and outcome, we summarized this variable five ways: as either a proportional or binary designation, with binary water cover value assigned based on either a 1%, 10%, 20% or 50% threshold. That is, given the 1% threshold, any cell with >1% water cover is considered to have water and any cell with >1% is not. For all species, we fit SDMs at all combinations of water cover variant, grain size, and season.

### Other model covariates

While our primary focus in this study was the influence of water cover on model performance and output, each SDM included several classes of covariates to account for numerous other factors that influence species distributions. Our topographic/habitat suite of covariates included mean elevation (from EarthEnv; Robinson et al., 2014), mean enhanced vegetation index (EVI; MODIS; https://lpdaac.usgs.gov/products/mod11a1v006/), topographic wetness index (TWI; hydroSHEDS; Marthews et al., 2015), and terrain roughness index (TRI; EarthEnv). Climatic covariates included mean annual temperature (bio1), mean annual precipitation (bio12), precipitation seasonality (bio15; all bio variables from CHELSA v2.1; Karger et al., 2021), and intra-annual variation in cloud cover (EarthEnv). Covariates were chosen from a pool of 32 candidates based on their minimal collinearity. Based on the mean value in each cell, all covariates that were not available at 1 km were spatially aggregated to 1 km. We also used temporal covariates in our models, such as year, date, and time of day, to account for temporal variability in bird activity and long-term population trends. Finally, as covariates, we included all effort indicators.

### Prediction surface

We first created a prediction surface with 1 x 1 km cells that covered our North American bounding box. All environmental covariates (topographic, climatic, and landcover suites) corresponding to each cell were generated at 1 x 1 km resolution. We masked portions of the prediction surface that fell outside of the buffered range extent when predicting the distribution of each species. We generated predictions for non-spatial covariates based on a 1 x 1 km, 1 hour search with 1 observer in 2020 on a randomized date within the season and at the hour of the day when the species is observed most frequently.

### Spatial grain

We mean-aggregated all spatial covariates assigned to each point (topographic, climatic, and percent landcover suites) and our prediction surface to the following spatial grain sizes (test scales) from 1 x 1 km to 5 x 5 km, 10 x 10 km, and 50 x 50 km to generate SDMs across spatial grains. The test grains were chosen based on their frequent use in published SDMs. Temporal and effort covariates were not scaled because they are assigned at the checklist (point) level.

### Species distribution models

We used Random Forest to model the breeding and non-breeding distributions of each species separately. Random Forest is a machine learning method designed to analyse large datasets with many covariates and is frequently found to produce the most accurate SDMs (e.g., Mi et al., 2017). Random forests are adaptable, automatically adjusting to complex, nonlinear relationships, and consider high-order interactions between environmental variables (Evans et al., 2011).

We randomly divided the data into training, testing, and out-of-bag (OOB) samples before modeling. The testing set was used for model validation, and the OOB set was used for threshold estimation. Using the ranger package (Wright et al., 2018), we fit five models for each species during each season corresponding to each test scale and water cover variant, resulting in 8,590 separate models. Models were parameterized to 100 trees and 7 threads. We compiled model predictive performance metrics including area under the ROC curve (AUC), true skill statistic (TSS), sensitivity, and specificity. By maximizing the sum of sensitivity and specificity, we determined optimal thresholds for each model. Following each model fit, we predicted to the prediction surface to generate thresholded grid-level occurrence predictions, calculating estimated range area for each model based on these predictions. To diagnose overfitting, we examined test-set calibration plots for each model.

## Results

As all performance metrics followed similar patterns, we focused on TSS for simplicity (see Figures A1, A2 for trends in AUC, sensitivity, specificity). In general, SDMs conducted at the finest resolution (1 x 1 km) performed very well (breeding TSS = 0.8; non-breeding TSS = 0.79; Figure 2). With coarsening resolution of environmental variables, model performance decreased. This drop in model performance was more pronounced for non-breeding season models. For example, while TSS dropped on average by 0.13 from 0.82 (1 x 1 km) to 0.7 (50 x 50 kilometres) for breeding season models, for non-breeding season models the drop was about 0.2. These patterns were independent of the summarization of water habitat variable (i.e., proportional versus binary; see Figures 2, A2).

**Figure 2.**
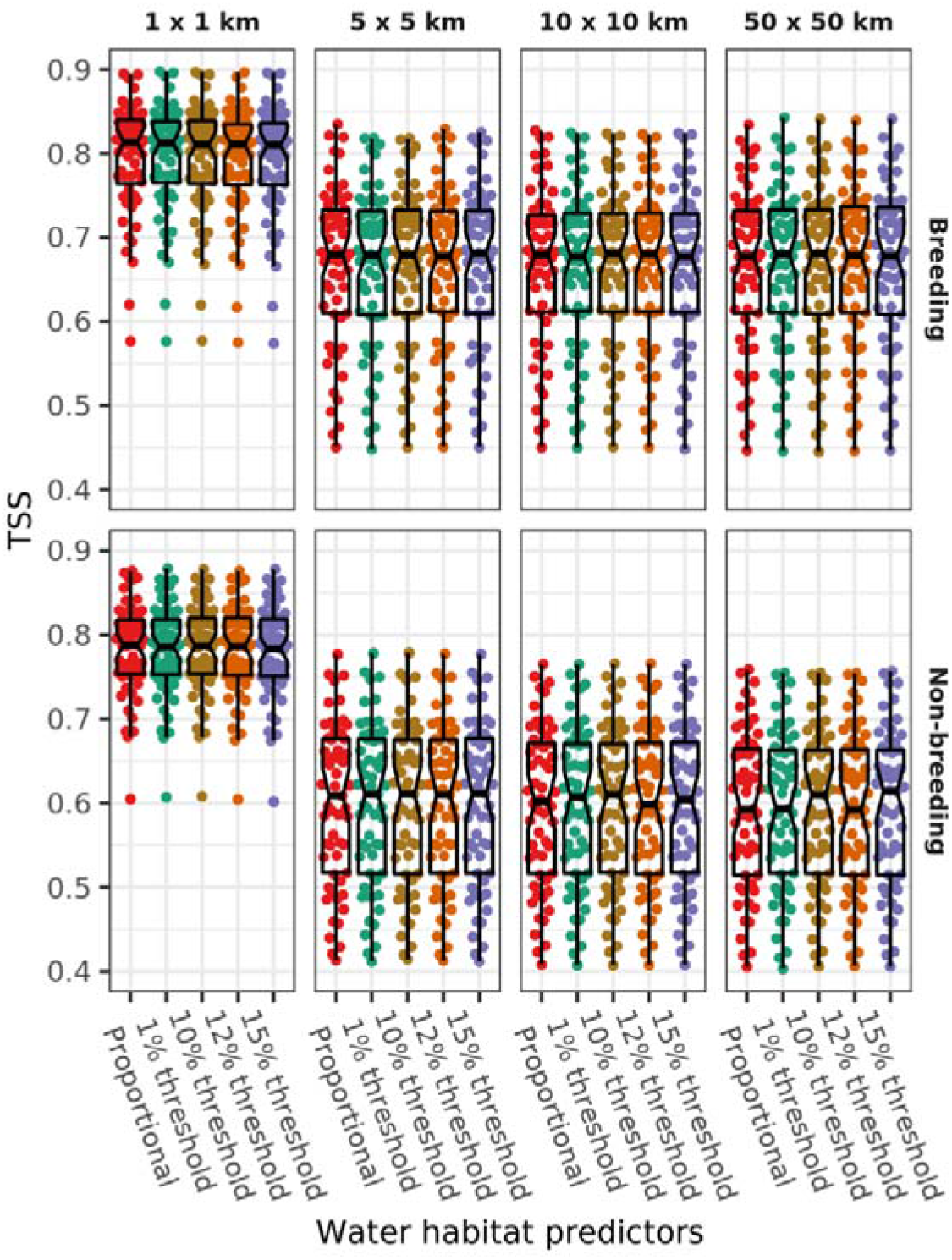
Predictive performance represented by TSS. Columns show results for different grain sizes, sorted by landcover summarization. Rows show results for different seasons. Boxplot lines represent median for each scenario, points represent individual species.

The comparison of model performance (TSS) between models fitted with water habitat as a proportional variable and as a binary variable revealed only minor differences in model performance (at 1 x 1 km resolution) or no differences (at 5 x 5 km and coarser resolutions) regardless of the threshold used to derive the binary water habitat variable (as seen in Figure 3A). Our results showed that the choice of landcover summarization mattered most in species with fewer observations, which are generally less common (Figure 3B). However, summarization of water cover did the influence predicted geographic range area for almost all (98%) species. We did not detect a particular bias – the median difference in predicted range was close to 0% (Figure 4). But for a good number of species models fitted with the binary variable over-or under-predicted the range area by more than 5%, and for some species this difference reached up to 30% (e.g., Cinnamon teal Spatula cyanoptera, see Figure 5). These patterns were consistent across spatial grains.

**Figure 3.**
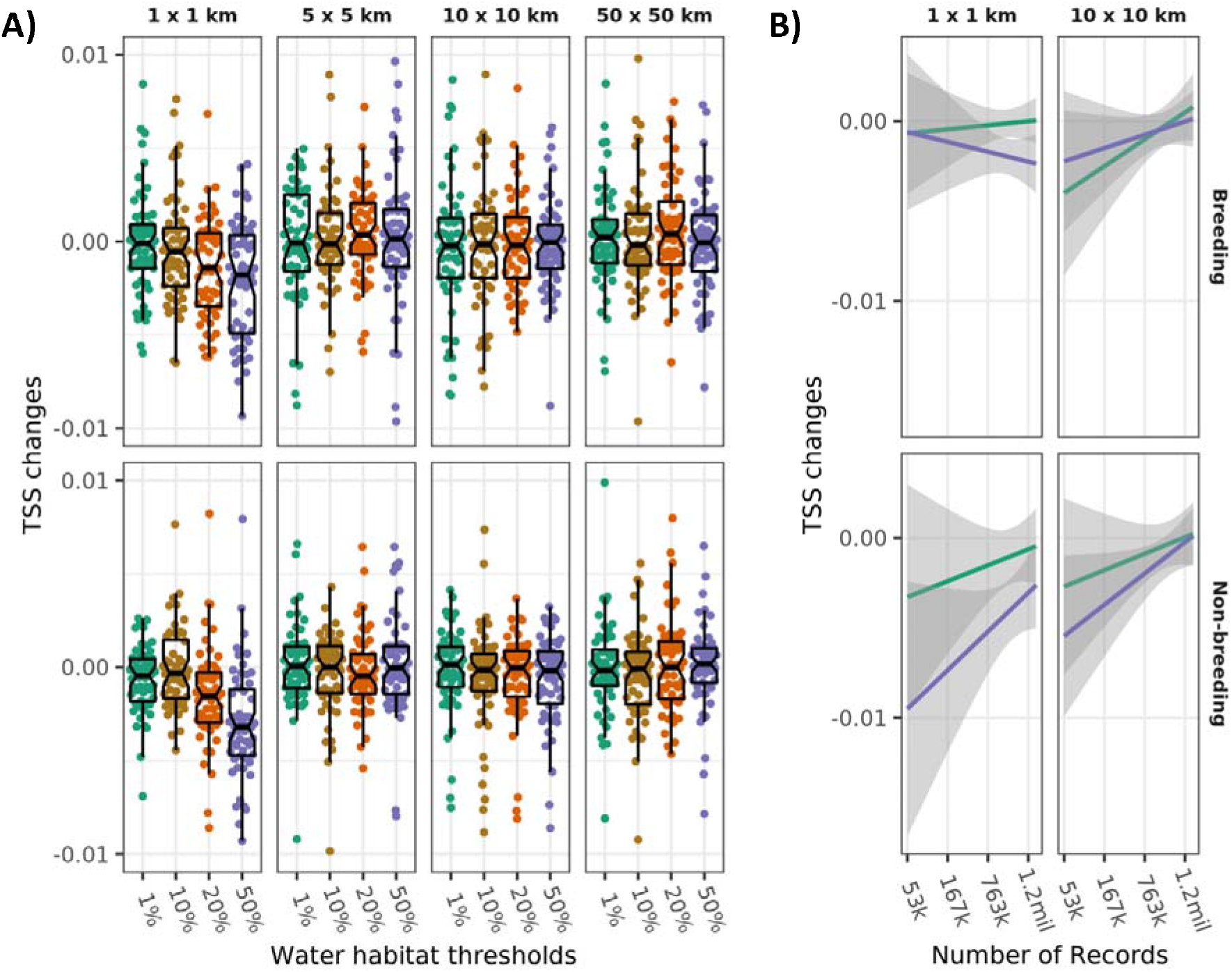
Variation of predictive performance across summarization method, sample size, and spatial grain. A: difference in TSS between models fitted with binary vs. proportional land cover. Boxplot lines represent median for each scenario, points represent individual species. B: variation in TSS across different sample sizes. Columns show results for different grain sizes. Rows show results for different seasons. Positive values indicate that models with binary habitat predictors performed better than those with proportional predictors and vice versa.

**Figure 4.**
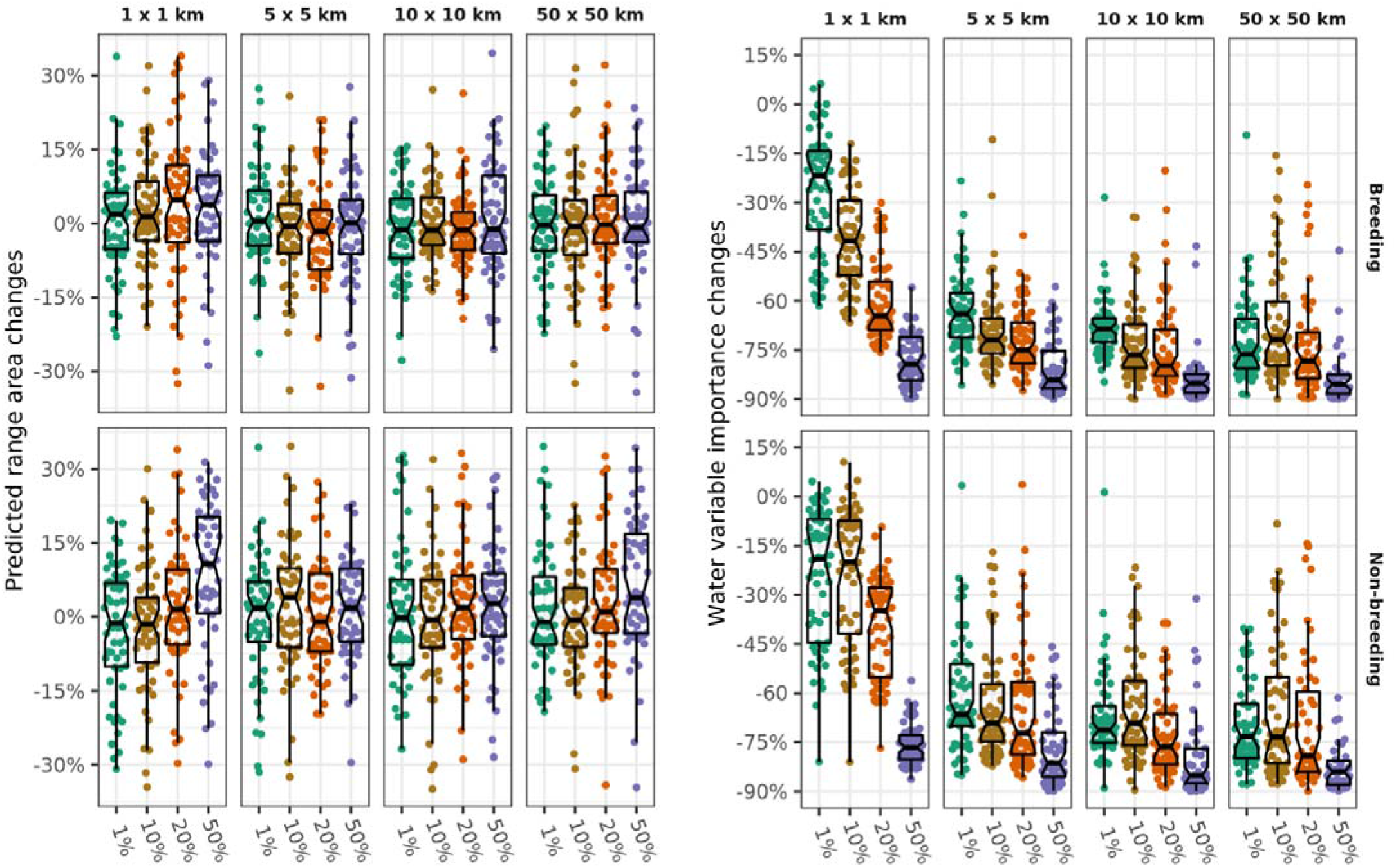
Differences in predicted range area (A) and variable importance (B) of models fitted with proportional vs. binary land cover (four thresholds). Columns show results for different grain sizes. Rows show results for different seasons. Boxplot lines represent median for each scenario, points represent individual species. Positive values indicate that models with binary habitat predictors predicted area respectively had higher water variable importance than models with proportional predictors and vice versa.

**Figure 5.**
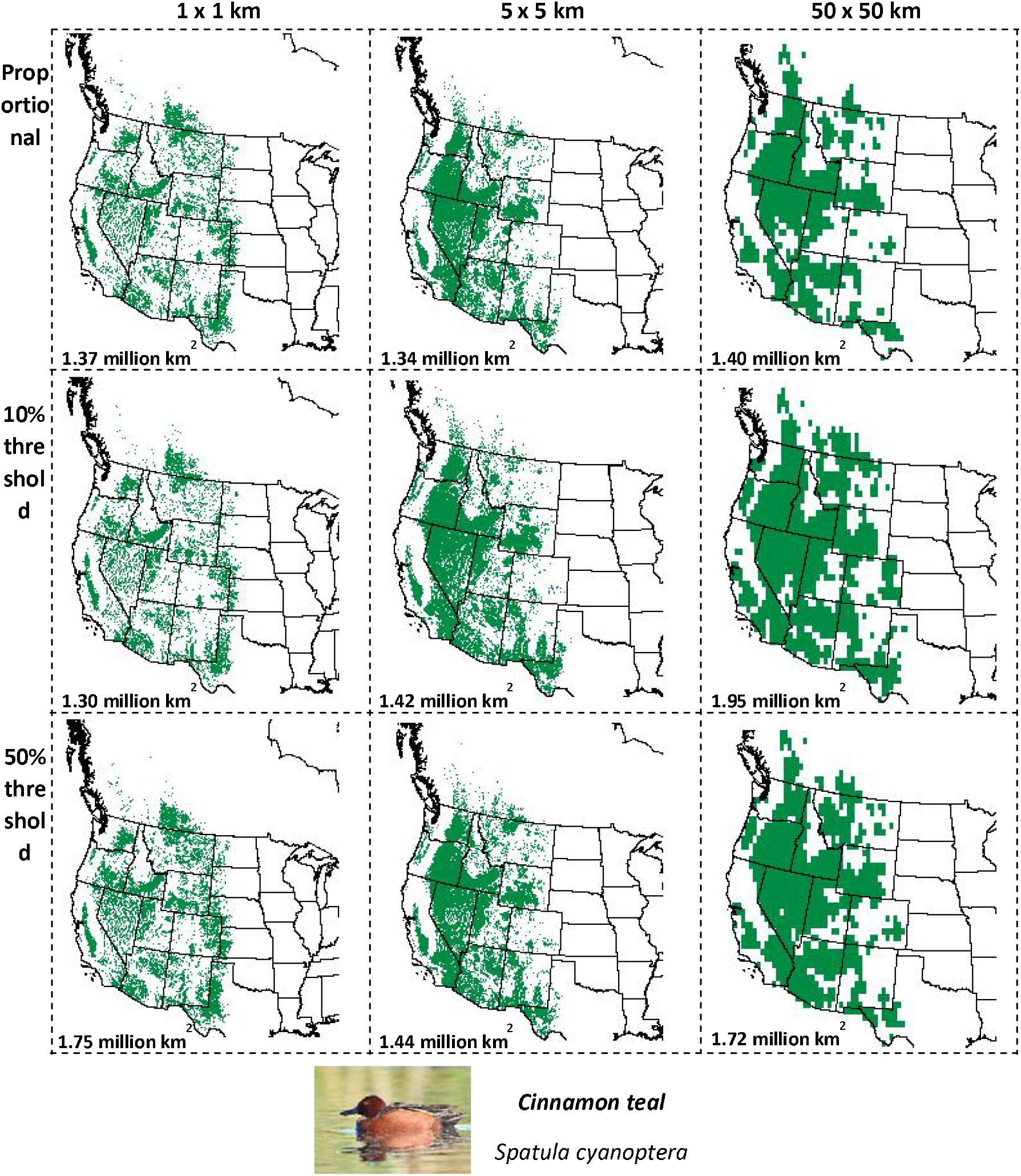
An example of changes in predicted range for Cinnamon teal (Spatula cyanoptera, n = 358, 252) between models fitted with binary and proportional land cover variables for non-breeding season.

Variable importance scores revealed that models with binary variables were relatively less informed by the water cover variable and instead other land cover features gained importance (Figure 6). For models fitted with binary variables, there was a significant decrease in the importance of the water variable (Figure 4). For models fitted with finest resolution of environmental variables (i.e., 1 x 1 km) the threshold played a role; the higher the threshold, the more pronounced the drop in water variable importance. For example, when models were fitted using a 1% threshold, the drop in variable importance was on average ca. 15%, whereas for models fitted using 50% threshold the drop in was over 70%. Toward coarser resolution of the variables, the differences between scenarios using various thresholds became smaller. At coarse resolution, all models fitted with binary water habitat variables significantly underestimated the importance of the water habitat variable (Figure 4).

**Figure 6.**
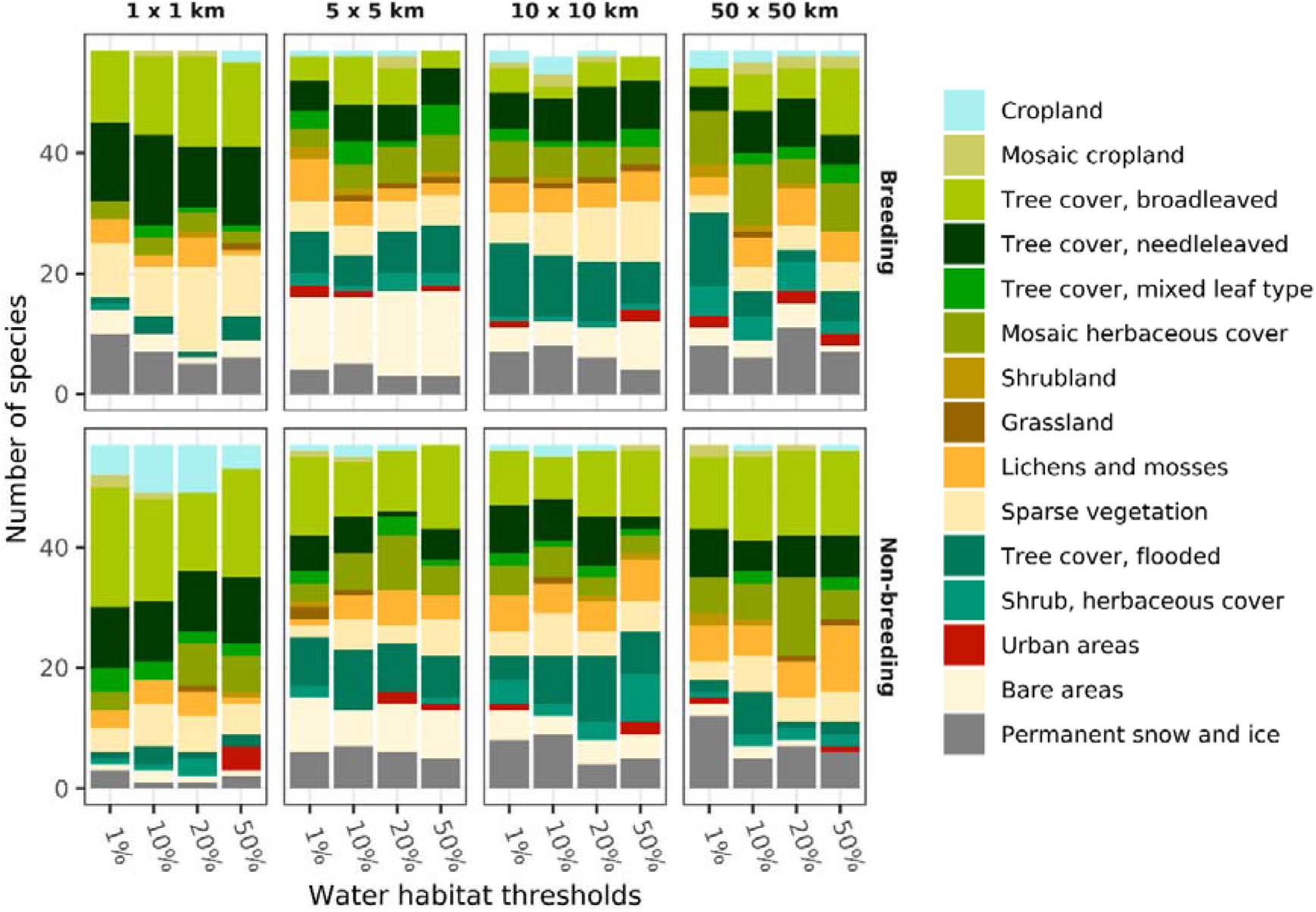
Habitat variables with the greatest increase in importance for random forest models fitted with binary variables compared with models fit with proportional variables.

## Discussion

Our study evaluated the usefulness of binary land cover variables for SDMs for species specialized in prevalent habitats. We developed species distribution models for 57 water bird species in North America across several grain sizes (i.e., from 1 km up to 50 km) for breeding and non-breeding seasons. We used proportional and binary land cover variables derived with various thresholds (1%, 10%, 20%, 50%). The results indicated that the performance of the models was not significantly affected by the type of the land cover variable used (proportional or binary), the season, or the threshold used to derive the binary variables (Figure 1A). This may be attributed to the large sample sizes used in the study, as North American birds are highly reported (the minimum number of occurrences per species was over 56,000 see Table A1). The largest changes in model performance were observed when using smaller sample sizes (Figure 3B), which is in line with previous studies that have explored the impact of sample size on SDMs (Stockwell and Peterson 2002, Wisz et al. 2008, Liu et al. 2019, Jiménez-Valverde 2020). Additionally, our study’s high number of occurrences helps ensure that the data represent most of the species realized niche, which can positively impact model performance regardless of the sample size (Boyd et al. 2022). In cases where a smaller sample size is used (tens or hundreds of records), a difference in model performance between models using proportional and binary variables would likely be more pronounced.

In contrast to model performance, using binary variables significantly impacted the models’ ability to detect water bodies as the most important land cover variable. We observed that with greater thresholds used to derive the binary water variable, the water cover became less important, with a drop of up to 90% at the coarsest grain used (see Figure 4). This drop can be explained by the coarse grain of environmental data (i.e., from 1 x 1 km up to 50 x 50 km). For example, a 1% threshold would require a water area equivalent to 247 American football fields at 1 x 1 km resolution, 25,000 at 10 x 10 km resolution, and 62,000 at 50 x 50 km resolution. For comparison, the area of the lake of the Ozarks (Missouri, USA) is only 54,000 football fields. Thus, it is apparent that even a small threshold can lead to a substantial drop in identified water bodies, which can still provide enough carrying capacity, food, and shelter for smaller water bird species populations (Hanski 1999, Melo et al. 2018). Our findings indicate that while binary variables can still lead to well-performing models with a low number of water bodies, other land cover variables may be identified as more important (Figure 6). For example, flooded tree cover or shrubland flooded tree cover or shrubland habitats often occur near water cover and may become predictive of a species occurrence when the species is also observed from these habitats (Figure 6), albeit less frequently, or species observations are imprecise in space.

The use of binary variables also influenced the predicted ranges of species across all spatial resolutions and seasons, regardless of the threshold used. While some predicted ranges varied significantly (e.g., the predicted range for Clark’s Grebe Aechmophorus clarkii was over 70% smaller when using a 10% threshold and 10 x 10 km spatial resolution), most models had over- and underprediction varying between 5% and 30%. These differences may result from models’ reliance on other land cover variables. They can have significant consequences for decision-making and conservation efforts, as they can result in high-priority conservation actions being directed at areas where the species do not occur and underestimate the importance of conservation actions in crucial areas for the species (Guisan et al. 2013, Velazco et al. 2020). If the predicted range area is underestimated, decision-makers may incorrectly design a protected area that is too small and that might be further disturbed by, for example, human activities. This can result in a population that is more vulnerable to factors such as predation, competition, inbreeding depression, or decreased likelihood of rescue effects after extinction events (Brown and Kodric-Brown 1977, Hanski 1999, Lande et al. 2003).

Our results indicate that as the grain size becomes coarser, the model performance decreases, which is consistent with prior studies (Guisan et al. 2007, Seo et al. 2009, Paradinas et al. 2022, Zarzo-Arias et al. 2022). Therefore, future studies should use the finest available grain size or the grain size at which species respond to the environment (Mertes and Jetz 2018, Gábor et al. 2022b). Importantly, our results suggest that the grain size can impact the applicability of binary land cover variables for models for species specializing in prevalent habitats. The smallest difference in the importance of water bodies between models fitted with proportional and binary variables derived using a 1% threshold suggests that in finer grain sizes, the hypothesis proposed by Gábor et al. (2022a) might be useful.

We modelled all species across both breeding and non-breeding seasons because we were unsure if the influence of landcover feature summarization was dependent on season. For example, we hypothesized that models with binary water cover perform worse when water is more critical to the biology of many birds during the breeding season, during which birds have offspring that often cannot leave the water or fly and many species remain stationary at a specific nesting site. Despite this, we found that the importance of water cover as a proportional feature is similar for both breeding and non-breeding seasons. This suggests that the method of land cover summarization is important even for organisms that are moving between habitats regularly, sometimes found in alternative habitat, or are otherwise not always tied to a specific habitat type.

As the world faces an unprecedented loss of biodiversity due to climate change and human activities, it is crucial to understand how and why species are distributed in space and time (Butchart et al. 2010, Barnosky et al. 2011, Jaureguiberry et al. 2022, Loreau et al. 2022). Advances in remote sensing and citizen science data have allowed ecologists to study these relationships on a scale and with a volume of species data like never before. To fully utilize this potential, we must prioritize understanding how to process and use these data effectively. This includes determining how to combine fine-resolution environmental data with species records of varying accuracy, finding the optimal grain size, and identifying the most accurate source of environmental data (e.g., Mertes and Jetz 2018, Moudrý et al. 2018, Lembrechts et al. 2019, Moudrý et al. 2019, Gábor et al. 2022b, Lu & Jetz 2023). As such, research can significantly improve the field of biodiversity conservation and help us understand the relationships between species and their environment. In our study, we contribute to this knowledge base by demonstrating that proportional data, compared to binary information, produce more accurate distributions for species specializing in prevalent habitat. However, we recommend further scrutiny of this finding for additional land cover types and for yet finers spatial grains (i.e., few or tens of meters).

## Data availability statement

Following our methods, bird occurrences and environmental data can be downloaded and processed as we did. R scripts to download data and develop species distribution models are available via repository (https://anonymous.4open.science/r/SDM-binary-land-cover-variables2).

